# Integration of Thermal Proteome Profiling with phosphoproteomic and transcriptomic data via mechanistic network models decodes the molecular response to PARP inhibition

**DOI:** 10.1101/2023.08.15.553354

**Authors:** Mira L Burtscher, Stephan Gade, Martin Garrido-Rodriguez, Ania Rutkowska, Thilo Werner, H Christian Eberl, Massimo Petretich, Natascha Knopf, Katharina Zirngibl, Paola Grandi, Giovanna Bergamini, Marcus Bantscheff, Maria Fälth-Savitski, Julio Saez-Rodriguez

## Abstract

The deregulation of complex diseases often spans multiple molecular processes. A multimodal functional characterization of these processes can shed light on the disease mechanisms and the effect of drugs. Thermal Proteome Profiling (TPP) is a mass-spectrometry based technique assessing changes in thermal protein stability that can serve as proxies of functional changes of the proteome. These unique insights of TPP can complement those obtained by other omics technologies. Here, we show how TPP can be integrated with phosphoproteomics and transcriptomics in a network-based approach using COSMOS, a framework for causal integration of multi-omics, to provide an integrated view of transcription factors, kinases and proteins with altered thermal stability. This allowed us to recover known mechanistic consequences of PARP inhibition in ovarian cancer cells on cell cycle and DNA damage response in detail and to uncover new insights into drug response mechanisms related to interferon and hippo signaling. We found that TPP complements the other omics data and allowed us to obtain a network model with higher coverage of the main underlying mechanisms. These results illustrate the added value of TPP, and more generally the power of network models to integrate the information provided by different omics technologies. We anticipate that this strategy can be used to broadly integrate functional proteomics with other omics to study complex molecular processes.

**Graphical abstract:** 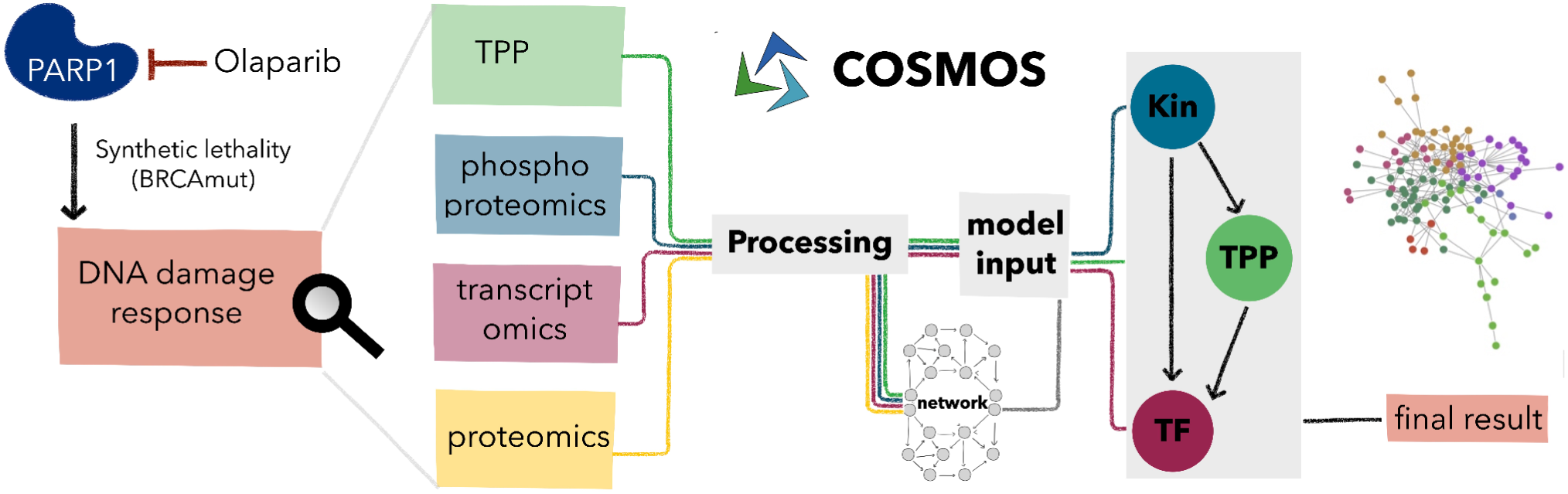

## 1. Introduction

Cellular regulation is a complex process mediated primarily by alteration in protein activity states and abundance. These alterations can be profiled via various omics technologies. Transcriptomics can be used to estimate the activity of transcription factors. Profiling post-translational modifications as in state of the art phosphoproteomics experiments is a way to explore protein function, in particular the activity of kinases. Thermal Proteome Profiling (TPP) ^1^ is a functional proteomics method that can identify shifts in protein thermal stability across different conditions ^2^. Changes in thermal protein stability can reflect a series of processes that influence protein structure and its interactions independent of changes in protein abundance ^3,4^. Especially in combination with quantitative changes measured by e.g. phosphoproteomics, TPP can be used to identify functionally relevant alterations and link them to signaling and phenotypic consequences ^1,2^. Due to the different strengths of these methods, an integrated study would not only shed light on their complementarities, but would allow us to explore their use in a synergistic manner.

To integrate the different layers of information encoded in each of these modalities, a series of computational methods have been developed, many of them making use of biological networks as integration scaffolds ^5^. Depending on the network content, different types of information can be leveraged. When networks are built based on previous discoveries, this existing prior knowledge can be explored systematically ^6^. COSMOS is a recently published method to integrate multi-omics data and prior knowledge to extract active sub-networks in a given functional context via causal reasoning ^7^. Within COSMOS, transcription factor and kinase activities are inferred from transcriptomics and phosphoproteomics data, respectively, and then causal paths are identified that connect the enzymes coherently with their inferred activities ^8^.

As a case study, we focused on the effect of PARP inhibition on DNA damage. Poly (ADP-ribose) polymerases (PARPs) are a family of 17 nucleoproteins, which transfer one or multiple ADP-ribose units (pADPr) to target proteins ^9^. This way, a series of cellular processes involving DNA repair, transcription, cell fate and stress response are regulated by PARylation ^10^. Upon PARP inhibition, DNA damage accumulates in cells initiating a cell cycle arrest and ultimately apoptosis, if the damage remains unrepaired ^11^. PARP inhibitors are used as drugs for targeted tumor therapy as they target cancer cells with distinct genetic defects in DNA repair genes specifically ^12^. Here, we use a TPP dataset acquired for ovarian cancer cells after incubation with a PARP inhibitor. We hypothesized that the integration of TPP data with other omics types will allow us to add context to the observed functional alterations and improve our mechanistic understanding of drug response. Clinical applications would benefit greatly from a better understanding of the differences between certain PARP inhibitors, the influence of patients’ genetic background, and potential resistance mechanisms ^12^.

Towards this aim, we integrated TPP, transcriptomics and phosphoproteomics data using the COSMOS framework in the context of PARP inhibition in ovarian cancer cells. We adapted COSMOS by modeling a signaling cascade comprising upstream kinases, intermediate proteins with altered thermal stability and downstream transcription factors (Figure 1). We found that a series of changes in thermal protein stability can be explained by deregulated kinase activities and phosphoproteomic changes. Further, we showed that the integration of TPP data is essential to model and extract known and novel pathways related to PARP inhibition in detail.

**Figure 1 -.**
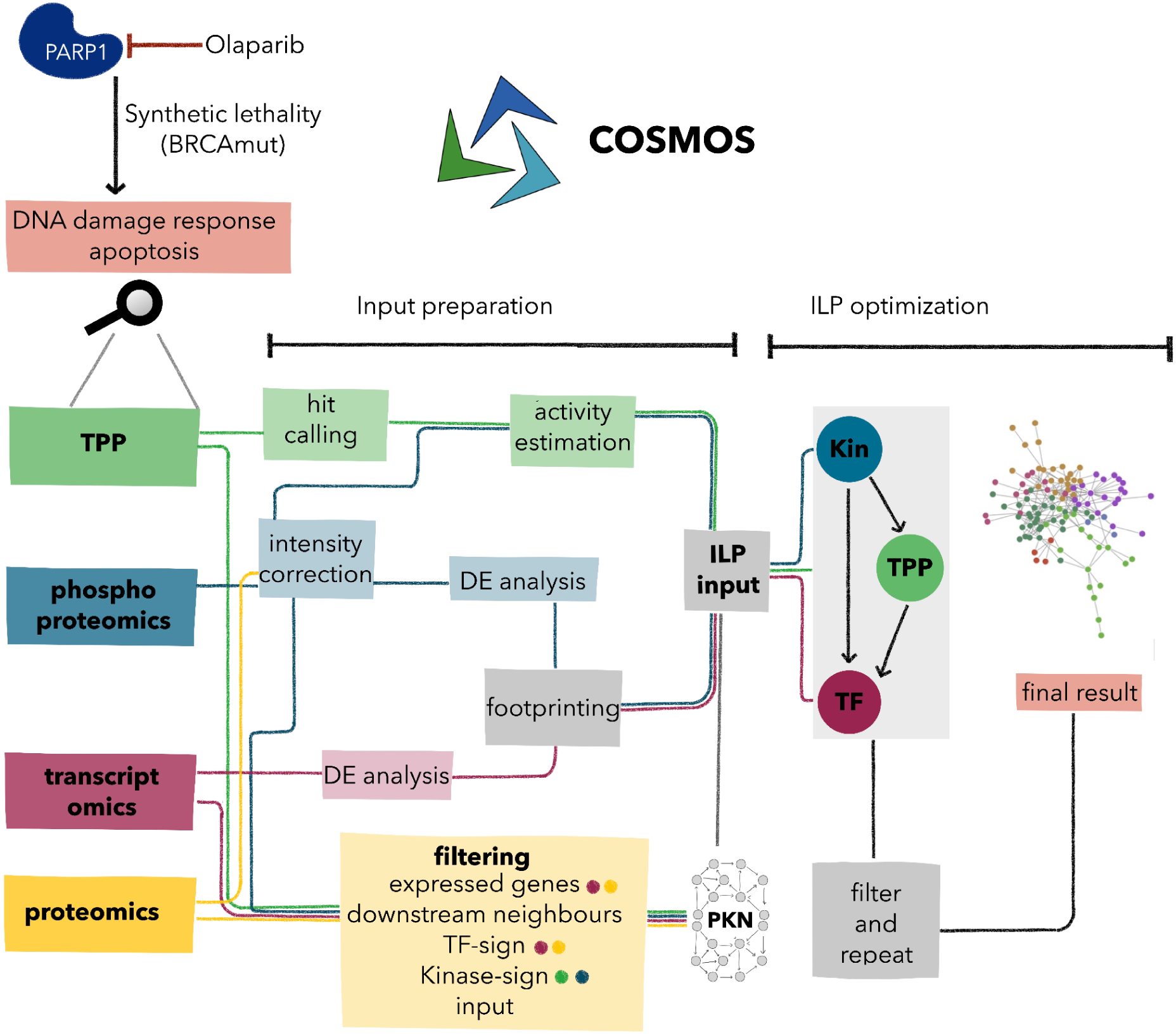
Overview of the COSMOS-TPP workflow. Preprocessing and modeling steps of the COSMOS-TPP workflow and the involved data types which are indicated by different colours. Olaparib is an inhibitor of PARP1, an essential player in DNA repair. Inhibition of PARP results in DNA damage that is not repaired in BRCA-mutated cells such as UWB1.289 ovarian cancer cells inducing apoptosis, a concept known as synthetic lethality. The different omic layers acquired after 24 hours of Olaparib treatment are processed separately and combined as input for an integer linear programming (ILP) optimization. Additionally, different omics information is used to filter the underlying prior-knowledge network (PKN) for example to only consider expressed genes in the final solution. To obtain a network model describing the interplay of Kinases (Kin), TPP hits and transcription factors (TF) two runs are merged to the final network. The coloured circles indicate the used data types in the respective filtering step.

## 2. Results

### 2.1 Multi-omic profiling of the response to Olaparib in ovarian cancer cells

To demonstrate the power of the proposed approach, we chose to investigate the effect of Olaparib in UWB1.289 cells. We characterized the response of UWB1.289 cells to Olaparib using transcriptomics, phosphoproteomics and TPP data (Supplementary Figure S1). In transcriptomics, we identified 20493 expressed genes, from which 44 changed significantly in response to the treatment (absolute logFC > log2(1.2) and adjusted P < 0.05). In phosphoproteomics, we identified 11615 phosphosites, 256 of which changed their abundance significantly in response to Olaparib (absolute logFC > log2(1.2) and adjusted P < 0.05). In the TPP data, we identified 9455 proteins, and found 76 that suffered thermal stability changes in response to the PARPi treatment (FDR < 0.1, Supplementary Table S3). In this case, a thermal (de)stabilization can correspond to a variety of effects ranging from drug binding to complex formation among others ^2^. In summary, we generated three independent datasets that inform about different types of molecular changes happening in response to Olaparib.

### 2.2 Footprint analyses reveal complementary molecular information between transcriptomics, phosphoproteomics and thermal proteome profiling

Next, we sought to explore if upstream regulators of the changes in transcriptomics and phosphoproteomics could offer complementary insight to the thermal stability changes detected through TPP. To this end, we performed a footprint analysis to estimate kinases and transcription factor activities from the phosphoproteomics and transcriptomics data, respectively (Supplementary Table 1, Supplementary Table 2). The analysis yielded activity estimates for 229 TFs and 170 kinases. The top 30 TFs showed a positive activity in response to the Olaparib treatment indicating a strong transcriptional activation (Figure 3A). Process-wise the top results comprised transcription factors involved in interferon (STATs, IRF1), nuclear receptor (RUNX1, ESR1), DNA repair (FOXM1, FOS/JUN) and cell cycle signaling (TFAP2C). For the phosphoproteomics data, the main deregulated kinases were found in the ATM-ATR axis which is known to mediate DNA damage checkpoint signaling (Figure 3B). Additionally, the downstream activation of numerous CDKs was also predicted by the footprint analysis, in line with the expected cell cycle arrest that occurs in response to PARP inhibitors. Overall, both transcription factors and kinases offered a molecular phenotype that explains the observed cellular response.

**Figure 2 -.**
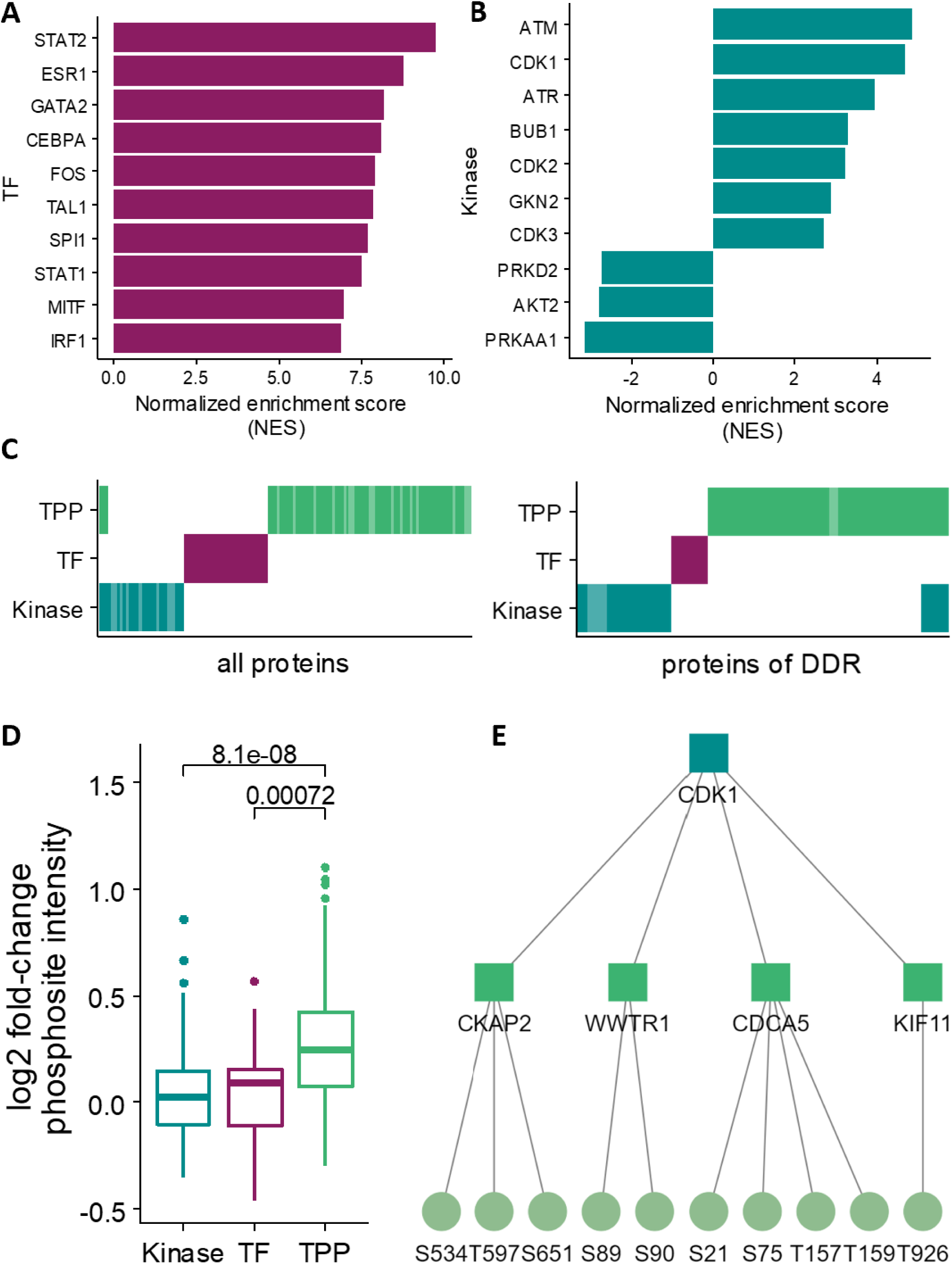
Transcriptomics, phosphoproteomics and thermal proteome profiling data exploration. (A) Normalized enrichment scores (NES) for top ten transcription factors and (B) top ten kinases as proxy of enzyme activity based on transcriptomics and phosphoproteomics data. In response to Olaparib treatment, cells react with a strong transcriptional activation and signaling around the DNA damage checkpoint (ATM, ATR, CDK1). (C) Comparison of top transcription factors, kinases and TPP hits overall and subsetted for proteins involved in DNA damage response. The different protein sets show almost no overlap, but functional complementarity to a certain extent as they all comprise several DDR members. The two overlapping proteins between the kinase and the TPP set are CDK1 and CDK2, both with a relevant role in cell cycle arrest upon DNA damage. (D) We compared log2 fold-changes of phosphosites on kinases, TFs and TPP proteins measured in the phosphoproteomics experiment. TPP proteins show a significant shift (p < 0.05) towards higher phosphosite intensities in comparison to Kinases and TFs. (E) TPP proteins further map frequently downstream of kinases identified in the activity estimation analysis as exemplarily shown for CDK1 with four downstream TPP proteins and related significant phosphosites.

**Figure 3 -.**
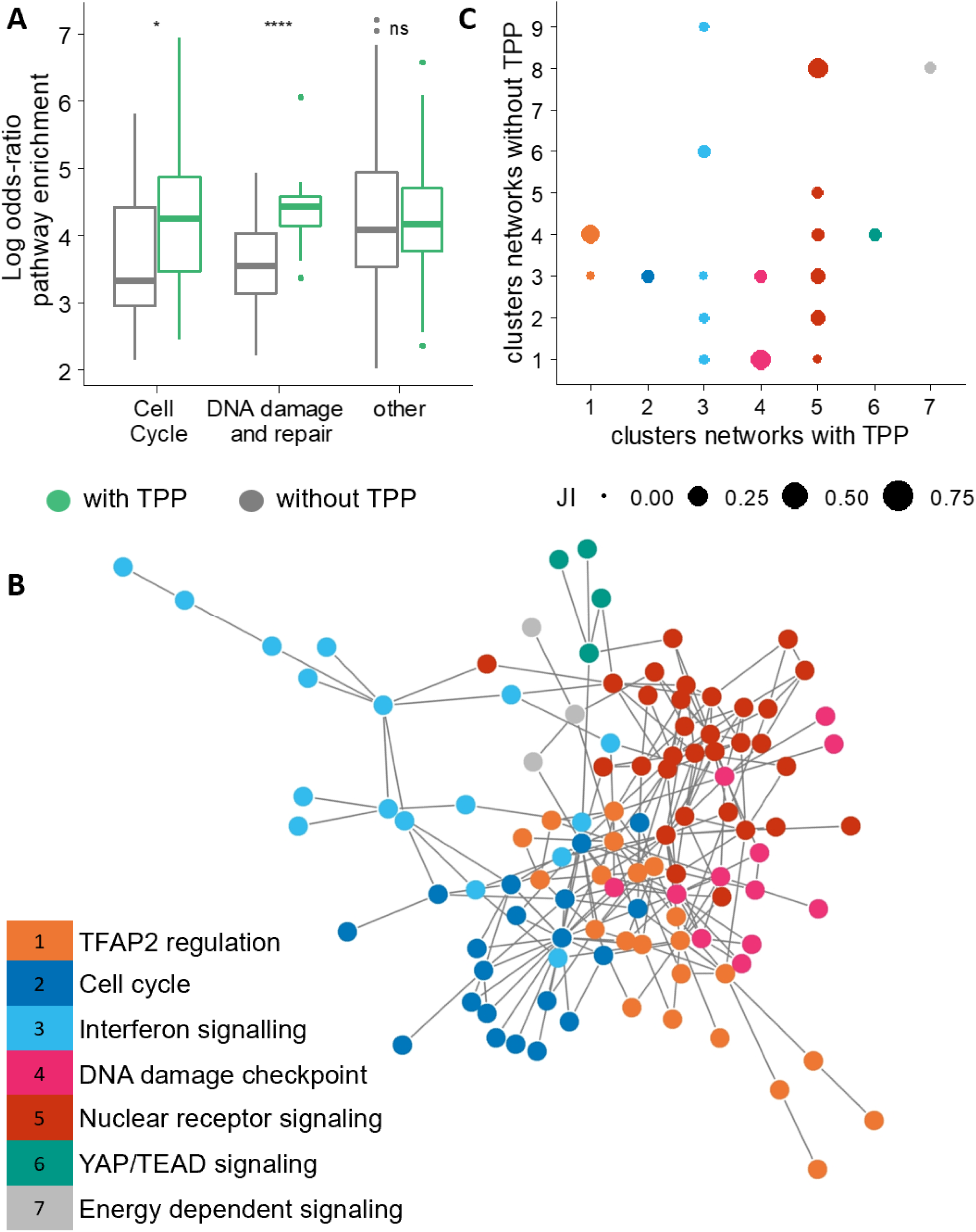
Effect of TPP integration on network level. (A) Comparison of the network-level enrichment results of networks with TPP data and networks without TPP. The enrichment results were filtered for significant pathways (q.value < 0.05, one-tailed Fisher’s exact test, Supplementary Table S5) and a minimum size of four nodes. We then extracted pathways related to cell cycle or DNA damage and repair and compared their log odds-ratio between networks with TPP and without TPP. (B) Graph representation of the network with TPP data. The colours indicate the seven clusters identified using a fast and greedy clustering algorithm. We performed a cluster specific enrichment of the nodes and summarized the significant pathways to one higher level process per cluster ranging from cell cycle to interferon signaling. (C) We compared the overlap of TPP and no TPP clusters using the jaccard index (JI). The colours refer to the respective cluster of the network with TPP. The size of the points refers to the jaccard index for the given comparison. The maximum overlap reached is a jaccard index of 0.28.

Regarding the thermal proteome profiling, the most prominent hits were CHEK2, PARP1, RNF146, MX1, different cyclins (stabilized) and RRN3 (destabilized). We found only a small overlap between TPP hits, transcription factors and kinases, both across all proteins but also in proteins belonging to the DNA damage response pathway (Figure 3C and 3D). Specifically, three proteins (CDK1, CDK2, CCNB1) overlapped between TPP hits and kinases, and we found no overlap between altered kinases and transcription factors. Despite the low overlap at the protein level, we found that the three omics layers inform about changes related to DNA damage response when we analyzed the pathways obtained from individual over-representation analyses.

Comparison of TPP data with other omics measurements revealed a high number of phosphorylations detected on TPP proteins (Supplementary Table S6). In detail, the TPP hits appeared to be more phosphorylated than kinases and transcription factors (Figure 1E), potentially indicating the relevance of the TPP hits in the signaling context. Furthermore, determining the number of direct upstream kinases of each TPP set revealed that 24 of the top 30 kinases are one step upstream to at least one TPP hit. One example of the connection between phosphoproteomics and TP is CDK1, with a direct edge to 20 TPP hits (Figure 1F). 55 phosphosites which were measured in the phosphoproteomics experiment can be mapped to these TPP proteins downstream of CDK1 (examples shown in Figure 3F).

All of this led us to hypothesize that the three data modalities offer complementary molecular perspectives on the same problem, and we therefore set out to apply an integrative approach to take advantage of the information they contain.

### 2.3 COSMOS robustly connects kinases and transcription factors with TPP hits

To integrate the three layers of molecular data we employed COSMOS ^7^, a recently developed method that integrates multi-omics data into causal molecular networks. To do so, it leverages the interactions that can be retrieved from prior knowledge databases, translating them into Integer Linear Programming constraints. Next, it solves an optimization problem that tries to extract the smallest possible network that causally connects the maximum amount of multi-omics measurements.

In total, we used 30 transcription factors, 30 kinases and 76 TPP measurements. We tested different approaches to integrate the three data layers, showing that more TPP hits can be included in the result if their activity is estimated beforehand, as in the case of kinases and transcription factors (Supplementary Figure S2A). As thermal stability changes do not inform about the sign of the deregulated activity (i.e., whether a protein activity is up or down regulated), we estimated TPP protein activity via phosphoproteomics data when possible. For the rest of the TPP proteins, we derived their activity during optimization and supported the activity inference with a (de-)stabilization weight (Supplementary Figure S2A). Regarding the network structure, we obtained the best results for networks using kinases as upstream regulators, TPP proteins as intermediate signaling proteins and transcription factors as downstream effectors (Supplementary Figure S2B). This COSMOS setup yielded a maximum of integrated TPP proteins in the result network.

Finally, we assessed the reproducibility and robustness of solutions (Supplementary Figure S2C). Analyses with different numbers of TPP proteins and different prior knowledge resources showed a good reproducibility of replicates, a robust clustering of networks with TPP versus networks without TPP and an expected influence of the prior knowledge resource. With this analysis we obtained a molecular network able to integrate and connect the information from the three omics layers.

### 2.4 Inclusion of TPP hits enhances biological information content in resulting networks

After ensuring the robustness and reproducibility of the integrated molecular networks, we aimed to profile the influence of TPP inclusion from a biological point of view. We performed an overrepresentation analysis of proteins in the networks with TPP or without TPP to identify processes which depend on the integration of TPP data to be modeled. The expected Reactome pathways around cell cycle and DNA damage were significant in both networks (q.value < 0.05, hypergeometric test) but significantly more prominent in networks with TPP (Figure 3A). Moreover, several of these DDR associated pathways such as mitotic checkpoint signaling, TP53 signaling, DNA damage recognition were significant in the networks with TPP exclusively. This again reinforces the complementary and synergistic aspect of the TPP with respect to the other two data modalities here analyzed.

To extract mechanistic insights in biological processes beyond the known and well characterized DDR, we performed a more granular cluster-based enrichment of the networks with and without TPP. The clustering of the network with TPP data resulted in seven clusters reflecting different biological processes (Figure 3C, Supplementary Table S4). Top pathways per cluster ranged from expected processes like cell cycle and DDR to new observations around interferon signaling and YAP/TEAD signaling. The later ones were not found in the enrichment analysis on a global level. In comparison, the clustering of the network without TPP yielded nine clusters with a very different composition (Figure 3B). The pairwise comparisons of clusters showed a maximum Jaccard Index of 0.28 for clusters related to DDR and nuclear receptor signaling. This cluster based network analysis allowed us to identify a series of PARP inhibition related processes such as DDR and interferon signaling and highlighted the value of TPP data to model these.

To investigate the role of TPP proteins in more granular biological processes, we analyzed three pathways identified in the cluster-based enrichment (Figure 3C). As a starting point, we focused on signaling around the DNA damage checkpoint and cell cycle arrest (Figure 4A). These processes are well-known to happen in response to PARP inhibition and depict the biggest part of the network ^11,13^. Besides the stabilization of PARP1 as a drug target, different proteins stabilized potentially upon phosphorylation as for example CHEK2, the major downstream target of the cell cycle arrest kinases, ATM and ATR ^14^. The network model also suggests the phosphorylation of multiple cyclins and cell cycle regulating proteins (CCNB1, CCNB2, CCNA2, BUB1B) by CDK1. At last, the TPP measurements also reflect complex formation as indicated exemplarily by the interaction of ESPL1 and PTTG1 which regulate the degradation of the APC/C complex during cell cycle arrest ^15^.

**Figure 4 -.**
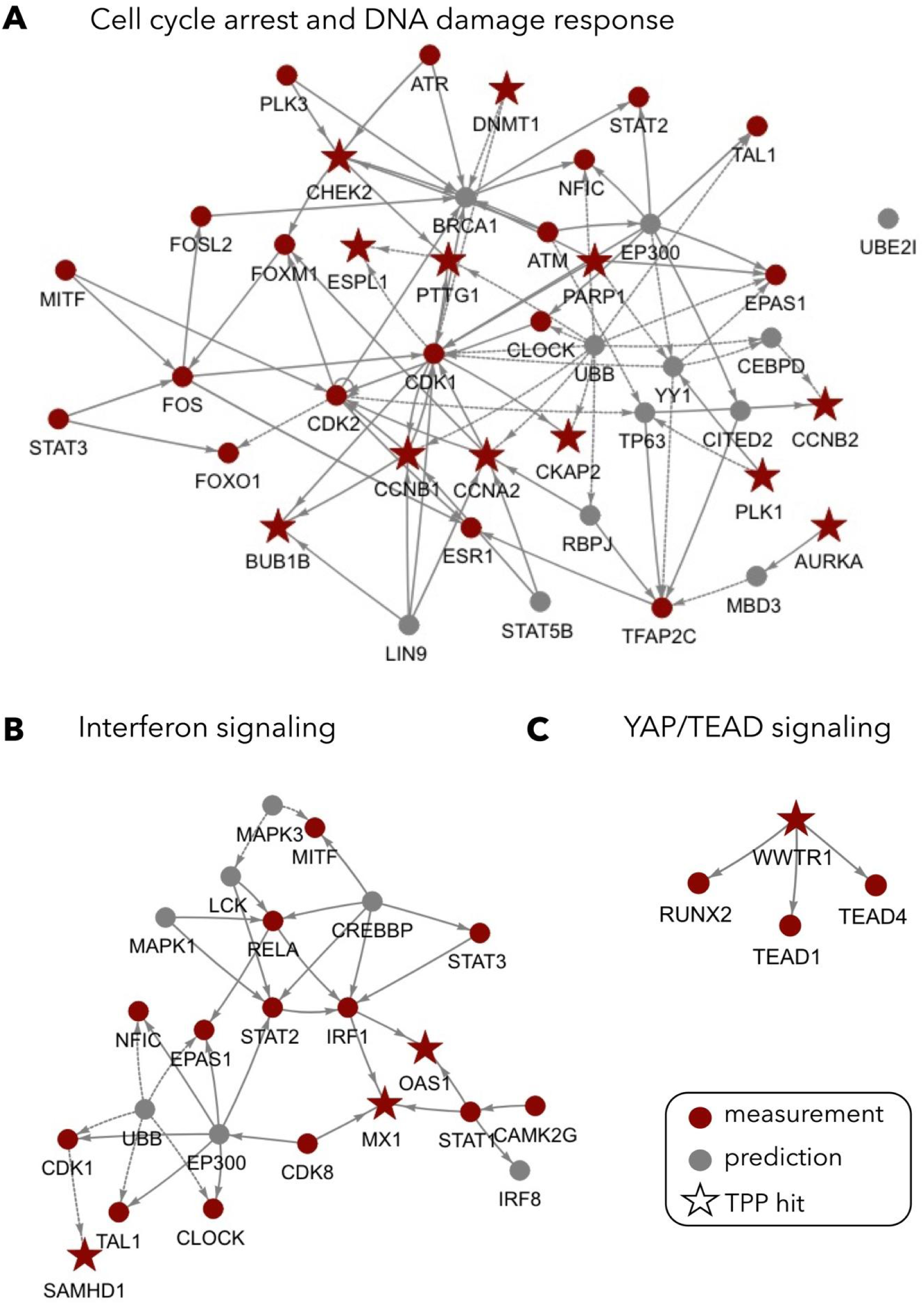
Mechanistic insight into selected molecular processes. Based on the results of the previous clustering analysis we extracted signaling related to the identified processes from networks with TPP to investigate it in detail. The red colour indicates TFs, kinases and TPP nodes which were used as input, while grey indicates predicted nodes added from the prior knowledge network during the optimization. TPP hits are indicated as stars. We chose three pathways represented by different clusters. (A) DNA damage response and cell cycle arrest, (B) Interferon signaling and (C) YAP/TEAD signaling.

Next we investigated interferon signaling, a known but less characterized response to PARP inhibition ^16,17^. Interferon signaling is represented around the transcription factors, RELA, STAT1, STAT2, STAT3 and IRF1 (Figure 4B). This transcriptional response is connected and complemented by the two TPP hits MX1, OAS1 which are activated upon interferon signaling ^18,19^. In this case, the thermal stabilization can be linked to transcriptional changes upon a stimulus.

Finally, we also investigated YAP-TEAD signaling, a not reported effect upon PARP inhibition (Figure 4C), but recently described to be central in DNA damage response ^20^. The complex of the TPP protein YAP and the TEAD1 and TEAD4 transcription factors forms in the nucleus at the end of the hippo signaling pathway ^21^. The subsequently initiated transcriptional program has been shown to contribute to an aggressive and drug resistant tumor phenotype ^22^.

## 3. Discussion

In this work, we integrated Thermal Proteome Profiling (TPP), phosphoproteomics and transcriptomics data from cancer cell lines subjected to drug-induced DNA damage using a causal-network approach. Overall, our results demonstrate that each individual omic profile carries a signal about the response of UWB1.289 cells to the PARP inhibitor Olaparib. A detailed footprint analysis showed that the three datasets could provide complementary insights into the molecular phenomena occurring after PARP inhibition, and we therefore created a molecular network that integrates them using our framework COSMOS ^7^.

PARP inhibitors represent a success story of modern precision medicine with numerous approved compounds for various cancers in less than 20 years after their discovery ^12^. Studies focusing on the effects of PARP inhibition at the transcriptome, proteome, or phosphoproteome level have been widely used to uncover synergies with other drugs, detect off-target effects, and characterize the pharmacological properties of various inhibitors, among others ^10,23–25^. Here we combine, within a network model, the information of these data modalities with thermal proteome profiling data. Our analysis helped us determine that the inclusion of TPP results contributed to 1) Capture known biological processes related to PARP inhibition and 2) Propose new hypotheses about lesser-known mechanisms of action.

TPP has been used successfully in the past for drug profiling, but it generates complex data that are difficult to interpret and potentially benefit from integration with other data types ^1,2,26^. By comparing the functional information provided by TPP with protein activity information derived from transcriptomics and phosphoproteomics data, we found that they complement each other in describing biological processes. The inferred transcription factors and kinases indicate the well-annotated players in the DNA damage response, whereas proteins with altered thermal stability reflect less annotated parts of the signaling space with strong links to the phosphoproteomic measurements which aligns with observations linking functionality of phosphosites with thermal stability ^3,27^.

To characterize such links and integrate the different data modalities, we used COSMOS ^7^. We show that we can integrate TPP data with transcriptomics and phosphoproteomics data and that this integration greatly improves the relevance of the resulting networks of COSMOS. At both the global and cluster level of the obtained networks, the inclusion of TPP enhances the reflected biological insights into PARP inhibition.

In particular, the inclusion of TPP allows us to retrieve detailed mechanistic insight into well-known consequences of PARP inhibition, such as DNA damage signaling via the ATM-ATR-CHEK2 axis or cell cycle arrest via modulation of cyclin degradation or the formation of the ESPL1-PTTG1 complex ^14,15^. In addition, we detect the induction of interferon signaling at the transcriptional level as well as less characterized effects such as signaling through the YAP1-TEAD complex indicated by a TPP and transcriptional signal ^16,17,20,21^.

This study is a first example of the potential of TPP in network-based multi-omics studies. More comprehensive data sets covering additional contexts and data modalities would be needed to draw conclusions about the generalizability of this approach. Given the advances in mass-spectrometry technology and throughput capacity, we anticipate that the number of studies generating functional proteomics datasets such as TPP will increase. We believe that generalizing this framework and extending it to additional data types such as solubility profiling or subcellular proteomics will be a useful step forward. Moreover, TPP data often contain more than just binary stabilization/destabilization information, such as effect size or distinct drug response curves, which could be used to improve modeling assumptions or downstream interpretation of the network ^28^.

In summary, we believe that the use of COSMOS to analyze TPP and other omics can help to extract mechanistic insight from these complex datasets.

## 4. Methods

### 4.1 Experimental procedures

#### Cell culture and Olaparib treatment

The UWB1.289 (CRL-2945; female) cell line was accessed from internal GSK collections and was obtained from ATCC. The cell line was additionally authenticated using the Promega Cell ID system. The generated short tandem repeat (STR) profiles matched exactly the expected STR profiles of the ATCC lines. Reagents were obtained from Gibco unless stated otherwise. Cells were cultured at 37°C, 5% CO_2_ in 1:1 RPMI:MEGM (Lonza, #CC-3151), 3% fetal bovine serum (FBS,Gibco), MEGM SingleQuots supplements (Lonza, #CC-4136) used without gentamycin-amphotericin. For the Olaparib (CAS: 763113-22-0) treatment, three 15 cm dishes per condition were prepared by seeding 3‒4 x 10^6^ cells per plate; 24 hours later medium was removed and 25 ml fresh medium containing dimethylsulfoxide (DMSO) (Sigma) or 4uM Olaparib was applied to the cells. Cells were incubated for indicated time (24 hours) at 37°C, 5% CO_2_. Cells were collected by trypsinization, washed twice with PBS, and counted using a Casy Cell Counter (OMNI Life Science). Cell pellets were generated containing 1–2 million cells for either proteomic and phosphoproteomic or transcriptomic analysis.

#### Transcriptomics

Approximately 1*10^6 UWB1.289 were lysed in 650 µl Tri reagent (Thermo Fisher, #AM9738) and bead milled with the following settings: 4 degrees Celsius, 2 cycles, 50% duty-cycle, 20 seconds, 4m/s. RNA was extracted using the RNeasy 96-well plate kit (Qiagen, #74181) according to manufacturer’s instructions including the DNase step. RNA concentration and integrity were assessed using a Fragment Analyzer (Agilent). Libraries were prepared using the NEBNext Ultra II Kit stranded (NEB, #E7760S) and NEBNext Poly(A) mRNA Magnetic Isolation Module (NEB, #E7490) following the manufacturer’s specifications using the following options: 600 ng of total RNA per sample (starting material), poly(A) enrichment (mRNA isolation), 8 PCR cycles, 10 min fragmentation time. DNA concentration and libraries size distribution were determined on a Fragment Analyzer. Libraries were pooled to 10 nM and sequenced on an Illumina NextSeq2000 instrument following the manufacturer’s specifications and aiming for approximately 40*10^6 reads per library. FastQ files were generated using the software bcl2fastq (version 2.20).

#### Proteomics and phosphoproteomics sample preparation

Cells were lysed in 4 % SDS, DNA was digested by benzonase following dilution to 1 % SDS. Lysates were cleared by centrifugation and the supernatant snap frozen until further processing. All samples were processed through a modified version of the single pot solid-phase sample preparation (SP3) protocol as described previously <u>29</u>. Briefly, proteins in 2% SDS were bound to paramagnetic beads (SeraMag Speed beads, GE Healthcare,#45152105050250,#651521050502) by addition of ethanol to a final concentration of 50%. Contaminants were removed by washing 4 times with 70% ethanol. Proteins were digested by resuspending in 0.1 mM HEPES (pH 8.5) containing TCEP, chloracetamide, trypsin and LysC following o/n incubation. Derived peptides were subjected to TMT labelling. The labeling reaction was performed in 100 mM HEPES (pH 8.5) 50 % Acetonitrile at 22 °C and quenched with hydroxylamine. Labeled peptide extracts were combined to a single sample per experiment.

For phosphopeptide enrichment, parallel samples were prepared as described above. Further sample preparation was performed on the BRAVO Assaymap (Agilent Technologies). Samples were desalted using C18-cartridges for AssayMap (Agilent Technologies) according to the software protocol provided by the manufacturer. In brief, samples were dissolved in 90 µl 6% TFA and loaded onto C18 columns equilibrated with 0.1% TFA. After loading, columns were washed with 0.1% TFA, followed by elution in 80% acetonitrile 0.1% TFA. For phosphopeptide enrichment, Fe(III)-cartridges for AssayMap (Agilent) were used on the BRAVO Assaymap (Agilent Technologies). Cartridges were primed with 50% ACN,0,1% TFA and equilibrated with 80% ACN, 0.1% TFA. 170µl 80% acetonitrile, 0.1% TFA were added to eluates from the C18 desalting step, and samples were loaded onto the Fe(III)-cartridges. Loaded Fe(III)-cartridges were washed with 80% acetonitrile, 0.1% TFA and phosphopeptides were eluted with 5% NH3 in water.

#### Thermal proteome profiling

Thermal proteome profiling was performed in live UWB1.289 cells in a 2D-TPP setting as described before ^1, 30^. In brief, cells were treated with 4 different Olaparib concentrations (0.4, 1, 4, 10 µM, or DMSO control) and incubated at 37 °C and 5% CO_2_ for 24 hours, then harvested by trypsinization and centrifugation. Cells were resuspended in PBS and transferred to 96-well polymerase chain reaction (PCR) plates. Cells were heated for 3 min to one of the 12 tested temperatures (42.1°C, 44.1°C, 46.2°C, 48.1°C, 50.4°C, 51.9°C, 54°C, 56.1°C, 58.2°C, 60.1°C, 62.4°C, 63.9°C). Cells were lysed with Igepal CA-630 0.8%, MgCl2 1.5 mM and benzonase 1 kU ml^−1^ and the aggregated proteins were removed by centrifugation through 0.45 µm filter plates. All flow-through from two adjacent temperature treatments were combined into a multiplexed TMT10 experiment. The database search was performed as described before ^1^.

#### Mass spectrometry (MS) analysis

All proteomic experiments utilized TMT for relative quantification. Measurements and analyses were performed as described before 31. TPPs and expression proteomics samples were off-line fractionated into 24 fractions, of which 8‒24 fractions were measured. Phosphopeptide enriched samples were measured without fractionation three consecutive times. Samples were measured on Thermo Orbitrap instruments (Orbitrap Lumos, Q Exactive HF, Q Exactive HFX, or Exploris). The database search was performed as described before 1; for phospho peptide enriched samples, phosphorylation of Ser/ Thr and phosphorylation of Tyr was further added as variable modification.

### 4.2 Data analysis

#### Preprocessing transcriptomic data

Raw FastQ files were processed to count matrices using the cloud processing tool DNAnexus. Adaptor trimming was carried out using Trimmomatic 32. Reads were then mapped to the reference human genome (GRCh38.96) with STAR v. 1.3.4 33. Picard MarkDuplicates v. 2.1.1 tool was used to identify and quantify PCR duplicates (http://broadinstitute.github.io/picard). Reads were assigned to genes using the command featureCounts from the software Subread to produce count matrices. Genes were prefiltered before differential expression testing to include only genes with more than 10 counts total across all samples. Statistics of differential gene expression were calculated with DESeq2 34. Resulting P values were adjusted for multiple testing. The criteria to consider a gene differentially regulated was an adjusted P value lower than 0.05 and an absolute log2 fold change in expression greater than 1.5.

#### Preprocessing proteomics data

Statistical analysis and visualization of the data were performed using the statistical language R. Phosphoproteomics data were aggregated to individual phosphosites. Log2 intensities were used as measure for phosphorylation and quantile normalised for sample to sample differences. Proteomics data were filtered for proteins with at least 2 unique quantified peptides. The log2 sum of ion intensities was used as a measure for protein abundance. These abundance measures were normalized using quantile normalization. Differential analysis was carried out using a moderated t-test implemented in the limma package ^35^. Resulting P values were adjusted for multiple testing. A protein was considered statistically significantly different with an adjusted P value below 0.05 and log2 fold change above log2(1.5) or below log2(1/1.5).

#### Footprinting

In order to estimate the activity of different input nodes (transcription factors (TFs), kinases, phosphatases) the viper algorithm was used ^36^. For transcription factors, the DOROTHEA database was used to obtain TF-target interactions with a confidence level of A, B or C ^37^. The interactions were downloaded using the OmnipathR package ^38^. For the viper algorithm, fold-change values of transcripts after limma analysis were used as input. The eset parameter was set to FALSE and the minimum target set size was set to 25 targets for one TF. Viper was further used to retrieve kinase/phosphatase activities. For this, normalized phosphosite intensities were corrected for total protein changes employing a linear model with the matched expression proteomics data. These corrected intensities were then used for differential testing before footprinting analyses. Phosphosite-enzyme collections were likewise downloaded with OmnipathR. Log2 fold-changes of phosphosites were used as input on phosphosite level. As for TFs, the eset parameter was set to FALSE and the minimum number of measured phosphosites per kinase was set to 3. Recently, a collection of methods to perform footprinting analyses in a comprehensive manner has been released ^39^.

#### TPP hit calling and activity estimation

To identify significantly (de)stabilized proteins as hits in the 2D-TPP dataset, we used a recently introduced method ^40^. In short, a null and an alternative model were fitted per protein and condition. The null model reflects a no-change hypothesis, while the alternative model describes concentration dependent protein abundance changes with temperature based constraints. Both models were fitted by minimizing the sum of squared residuals and compared using an F-statistic. Significance was adjusted by a bootstrap-based false-discovery rate (FDR) procedure. We evaluated the TPP hits by comparison with the other omic data types and a pathway enrichment analysis. We performed a TPP activity estimation based on causal prior knowledge links between kinases and TPP proteins. TPP hits were compared to direct upstream kinases which were also among the top 30 regulators identified in the footprinting analysis. If the activity of all upstream kinases of one TPP hit and their prior knowledge link to the TPP hit were consistent, the TPP activity was inferred from this (i.e. Kinase A is active and has an activation PKN link leading to a TPP protein, it is active as a consequence). Further, links between kinases and TPP phosphorylation sites which were not in agreement were removed from the PKN in a Kinase-sign filtering step. For the COSMOS robustness analysis, additional TPP hit sets were generated using a strict approach evaluating curve fit (R2 > 0.8 with sigmoidal model) and effect size (logFC > 1.5) as well as the top 100 proteins according to a stabilization score introduced by ^26^.

#### Pathway analysis

For all enrichment analyses we used the ReactomePA package 41. For the protein activity analysis results we used the top 30 regulators as input, for the TPP hit calling results all hits detected and on network level all nodes of the result network. As background we chose all genes, phosphosites or proteins detected for the single omics datasets and all nodes from the prior knowledge network for the network-level enrichments. In the enrichPathway() function the organism was set to “human”, minGSSize was set to five and maxGSSize was set to 500. To identify significant pathways we filtered the result with a significance cutoff of 0.05 after multiple testing correction.

#### CARNIVAL/COSMOS and prior knowledge resources

To infer signaling processes, the CARNIVAL algorithm combines upstream perturbation information (i.e. drug targets) with downstream measurements (i.e. transcription factor (TF) activities) and a causal prior knowledge network (PKN) ^42^. The enzyme activities are calculated using a footprinting method ^8^. The PKN contains the signed and directed protein-protein interactions (node A activates/inactivates node B) which are used to model signaling interactions between measurements and perturbation input. The perturbation and measurement input as well as the PKN are then used to minimize an objective function in an integer linear programming (ILP) optimization. In COSMOS, upstream kinases, phosphatases and TFs are connected with downstream metabolites expanding the use of CARNIVAL to a multi-omics context ^7^. In this study, we used a prior knowledge network which is provided as part of the COSMOS package (30k interactions) and a second prior knowledge network based on protein-protein interactions of metabase (117k interactions). The prior knowledge resources were filtered and adapted as described by Dugourd et. al ^7^. As measured input we used the top 30 transcription factors and kinases/phosphatases from the activity estimation analysis as well as the 76 hits of the TPP dataset. Eleven of these TPP hits had an activity estimate, the others were used with a weight reflecting (de)stabilization.

#### Setup of COSMOS with TPP

We first used kinases and TPP proteins as upstream nodes and transcription factors as downstream nodes. This represents the signaling cascade in response to a stimulus going through kinases and proteins with altered thermal stability before it causes transcriptional adaptation. The prior knowledge then was filtered in between runs as described by Dugourd et. al. before we performed a second run from kinases upstream to TPP proteins and transcription factors downstream to complete the network. Both runs were repeated once to improve the result quality. We made the union of the two final networks, resulting in a combined complete network. To test whether CARNIVAL’s formulation in combination with the adaptations for multi-omics introduced with COSMOS can be used this way, we tested its reproducibility in different settings. We generated a bigger and a smaller TPP protein set (100 and 23 hits) using alternative hit calling strategies ^26^. For all three TPP sets as well as kinases and transcription factors alone (noTPP), we produced three network replicates for each of the two prior knowledge resources. We compared the results based on the integration of nodes and their activity prediction.

#### Network enrichment analysis and comparison

For the three replicates of the network with TPP hits determined using the F-statistic based results and three network replicates without TPP, we performed a node enrichment analysis as described using ReactomePA ^41^. We compared the results of the enrichments by calculating the log odds ratio for each pathway and extracting pathways relevant for cell cycle and DNA damage response. A t-test was used to determine the differences between biological processes in networks with TPP and networks without TPP.

#### Clustering, cluster comparison and cluster enrichment

To retrieve a more granular view of biological processes on network level we performed a clustering analysis. We extracted all edges identified in one replicate for networks with F-statistic based TPP hits and networks without TPP hits, respectively and applied a fast and greedy clustering algorithm on the resulting graph. The pairwise Jaccard index was used to compare the composition of the resulting clusters of networks with TPP and without TPP. We performed a cluster-specific enrichment analysis as described before and extracted all nodes related to the identified processes to gain mechanistic insight.

## Code availability

All code used to perform the computational analyses described is available at https://github.com/saezlab/COSMOS-TPP.

## Data availability

All raw files, search parameters and search outputs were deposited to the ProteomeXchange Consortium via the PRIDE partner repository and will be made accessible upon publication. RNA-seq data have been deposited in the Gene Expression Omnibus (GEO) databank and will be made accessible upon publication.

## 5 Acknowledgements

We thank Matthias Kalxdorf, Nao Iwamoto and Aurelien Duguord for very helpful discussions and analysis support during this project. This publication was supported through state funds approved by the State Parliament of Baden-Württemberg for the Innovation Campus Health + Life Science Alliance Heidelberg Mannheim.

## 7. Authors contributions

Burtscher M.: Conceptualization, Software, Visualisation, Investigation, Interpretation, Writing-Original draft preparation. Gade S.: Conceptualization, Software, Supervision, Writing-Original draft preparation. Garrido-Rodriguez M.: Writing-Original draft preparation, Writing - Review & Editing. Rutkowska-Klute A.: Investigation, Data curation. Eberl H.C: Investigation, Data curation. Werner T.: Investigation, Data curation. Petretich M.: Investigation, Data curation. Knopf N: Investigation, Data curation. Zirngibl K.: Conceptualization. Grandi, P.: Project administration, Supervision. Bergamini G.: Project administration, Supervision. Bantscheff M.: Project administration, Supervision. Faelth-Savitski M.: Project administration, Supervision, Writing - Review & Editing. Saez-Rodriguez J.: Conceptualization, Project administration, Supervision, Writing - Review & Editing.

## 8. Supplementary Materials

### Tables

All supplementary tables can be found in the provided excel file.

**Supplementary table 1. Kinase enrichment results**

**Supplementary table 2. Transcription factor enrichment results**

**Supplementary table 3. TPP hits and scores**

**Supplementary table 4. Network clustering results**

**Supplementary table 5. Pathway enrichment results**

**Supplementary table 6. TPP phosphorylation**

### Figures

**Supplementary Figure S1 -.**
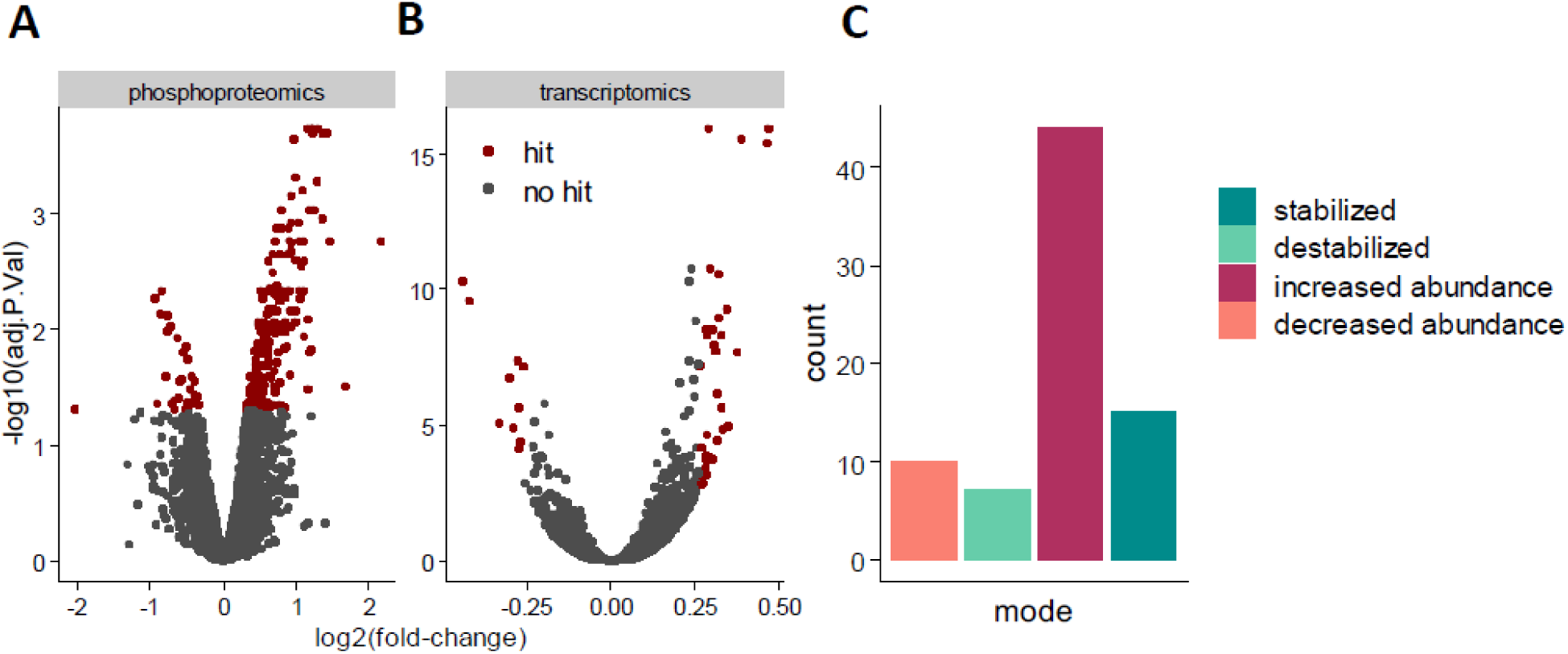
Transcriptomics, phosphoproteomics and TPP data. (A) Volcano plot of the phosphoproteomics data after limma analysis comparing 4 uM Olaparib vs DMSO control. (B) Volcano plot of the transcriptomics data after limma analysis comparing 4 uM Olaparib vs DMSO control. (C) Overview of the TPP hits determined using the F-statistic based hit calling approach comparing the number of (destabilized) and abundance changed hits.

**Supplementary Figure S2 -.**
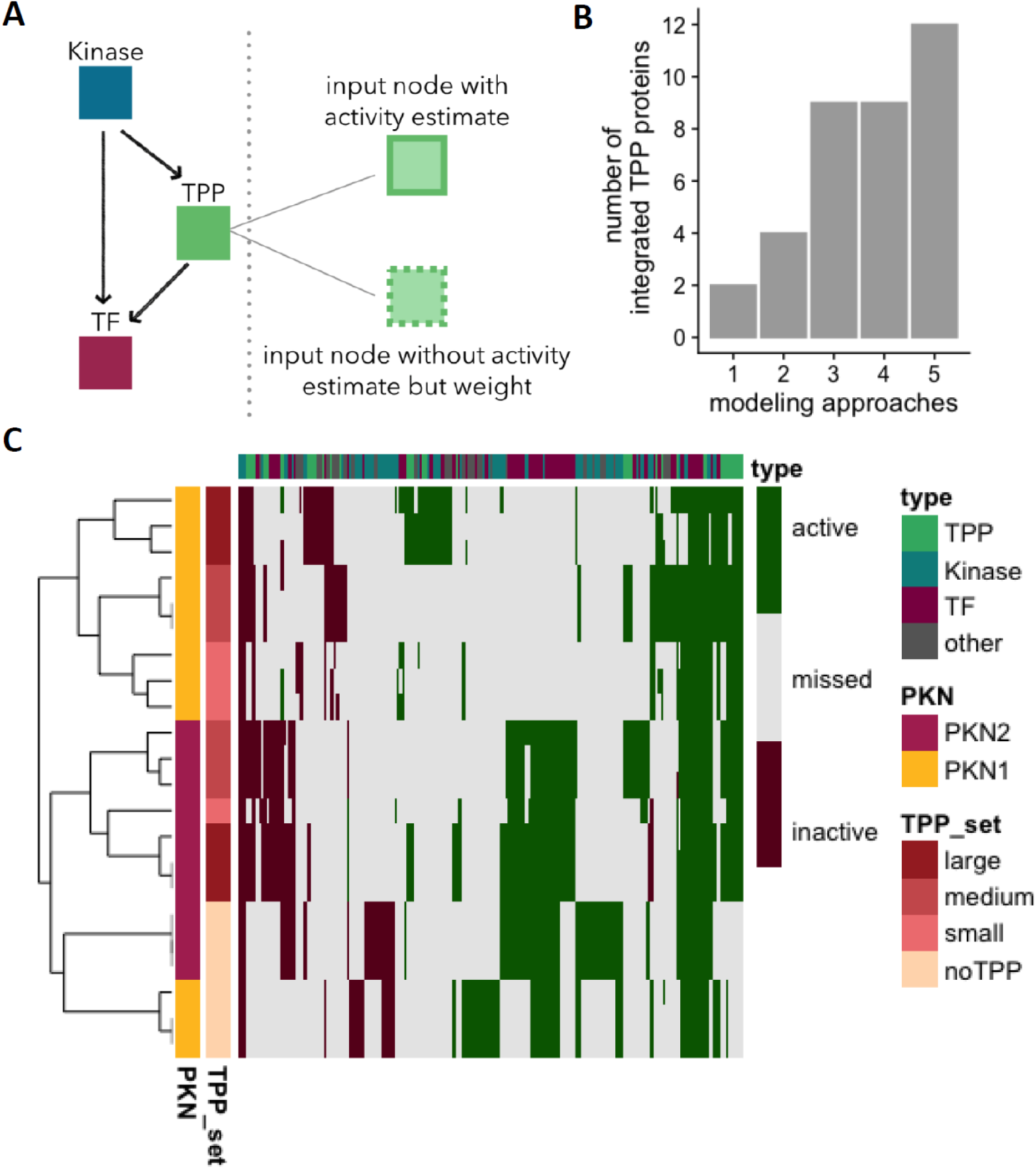
Robustness and reproducibility. We tried to validate whether the chosen setup (combination of runs, activity estimation and filtering steps) can be used to model reasonable networks with COSMOS. In principle, the integer linear programming solver optimizes towards a local optimum and the complete search space of the problem is not known. As a consequence, multiple technical replicates of COSMOS can have similar but slightly different optimal solutions. To assess if this solver-dependent variability can be distinguished from actual differences due to the used input data, we set up a small robustness analysis. (A) We integrated TPP proteins between upstream kinases and downstream transcription factors into the signaling cascade based on the correlation of phosphoproteomic and TPP data. Further, we implemented a strategy for activity estimation where we inferred activities for TPP proteins via upstream kinases and phosphorylation status. For TPP proteins without activity estimate we inferred the activity state during the optimization supported by a weight reflecting (de)stabilization. Different modeling setups (1-5) were tested to maximize the number of integrated TPP proteins which we reached using activity estimations and weights. Details of the optimization process are provided in the CARNIVAL and COSMOS publications ^7,42^. (B) We clustered replicate networks for different TPP set sizes based on node activities (active, missed, inactive). All replicate runs of distinct TPP sets cluster together. The networks separate into networks with TPP input and networks without TPP input. Further the influence of the used prior knowledge is clearly visible (PKN).

|  |  |
| --- | --- |
| 1 | no TPP proteins as input |
| 2 | TPP proteins without activity information |
| 3 | TPP with activity estimation |
| 4 | TPP with activity estimation and weights |
| 5 | TPP with activity estimation, NA values and weights |

